# Modular assembly of the nucleolar large subunit processome

**DOI:** 10.1101/223412

**Authors:** Zahra Assur Sanghai, Linamarie Miller, Kelly R. Molloy, Jonas Barandun, Mirjam Hunziker, Malik Chaker-Margot, Junjie Wang, Brian T. Chait, Sebastian Klinge

## Abstract

Early co-transcriptional events of eukaryotic ribosome assembly result in the formation of the small and large subunit processomes. We have determined cryo-EM reconstructions of the nucleolar large subunit processome in different conformational states at resolutions up to 3.4 Ångstroms. These structures reveal how steric hindrance and molecular mimicry are used to prevent premature folding states and binding of later factors. This is accomplished by the concerted activity of 21 ribosome assembly factors that stabilize and remodel pre-ribosomal RNA and ribosomal proteins. Mutually exclusive conformations of these particles suggest that the formation of the polypeptide exit tunnel is achieved through different folding pathways during subsequent stages of ribosome assembly.

## Introduction

The assembly of the large eukaryotic ribosomal subunit (60S) is organized as a series of consecutive intermediates, which are regulated by different sets of ribosome assembly factors in the nucleolus, nucleus, and cytoplasm (1). During the past decade, three major types of pre-60S particles have been observed, each with a distinct composition of assembly factors, representing late nucleolar (2), nuclear (3) and late nucleo-cy-toplasmic intermediates (4). The earliest pre-60S particles exist in the nucleolus, where they undergo major RNA processing steps and conformational changes. Subsequently, in the nucleoplasm, the internal transcribed spacer 2 (ITS2) RNA is removed and the particles are assembled into a nuclear export competent state. Exported particles then complete the final steps of maturation in the cytoplasm.

While previous high-resolution structural studies have revealed the architectures of the late nucleolar and export-competent pre-60S particles, the early nucleolar particles, containing specific factors such as Nsa1 have so far remained elusive (2, 4). Although the identities and approximate binding regions of many early nucleolar ribosome assembly factors are known, their structures and functions have not yet been determined (5).

## Structure of the LSU Processome

To elucidate the mechanisms that govern early nucleolar large subunit assembly, several labs, including ours have observed that cellular starvation elongates the lifetime of a 27SB pre-rRNA containing species (6, 7). Here, we have isolated these intermediates from starved yeast by employing a tandem affinity purification involving tagged ribosome assembly factors Nsa1 and Nop2 (**fig. S1**). While compositionally related to Nsa1-containing particles (8), we refer to this nucleolar pre-60S particle as the large subunit (LSU) processome, as it accumulates upon starvation like the small subunit processome. Purified LSU processomes were analyzed by cryo-EM leading to the discovery of three states, which were determined at resolutions of 3.4 Å (state 1), 3.7 Å (state 2) and 4.6 Å (state 3) (**fig. S2, S3, table S1**). These three states, together with cross-linking and mass spectrometry data, resulted in the identification of 21 ribosome assembly factors (table S2 and data table S1). Atomic models could be built for a large number of these proteins (**fig. S4**), while homology and poly-alanine models were used to build proteins located in more flexible regions (**Fig. 1, fig. S4**).

**Fig. 1.**
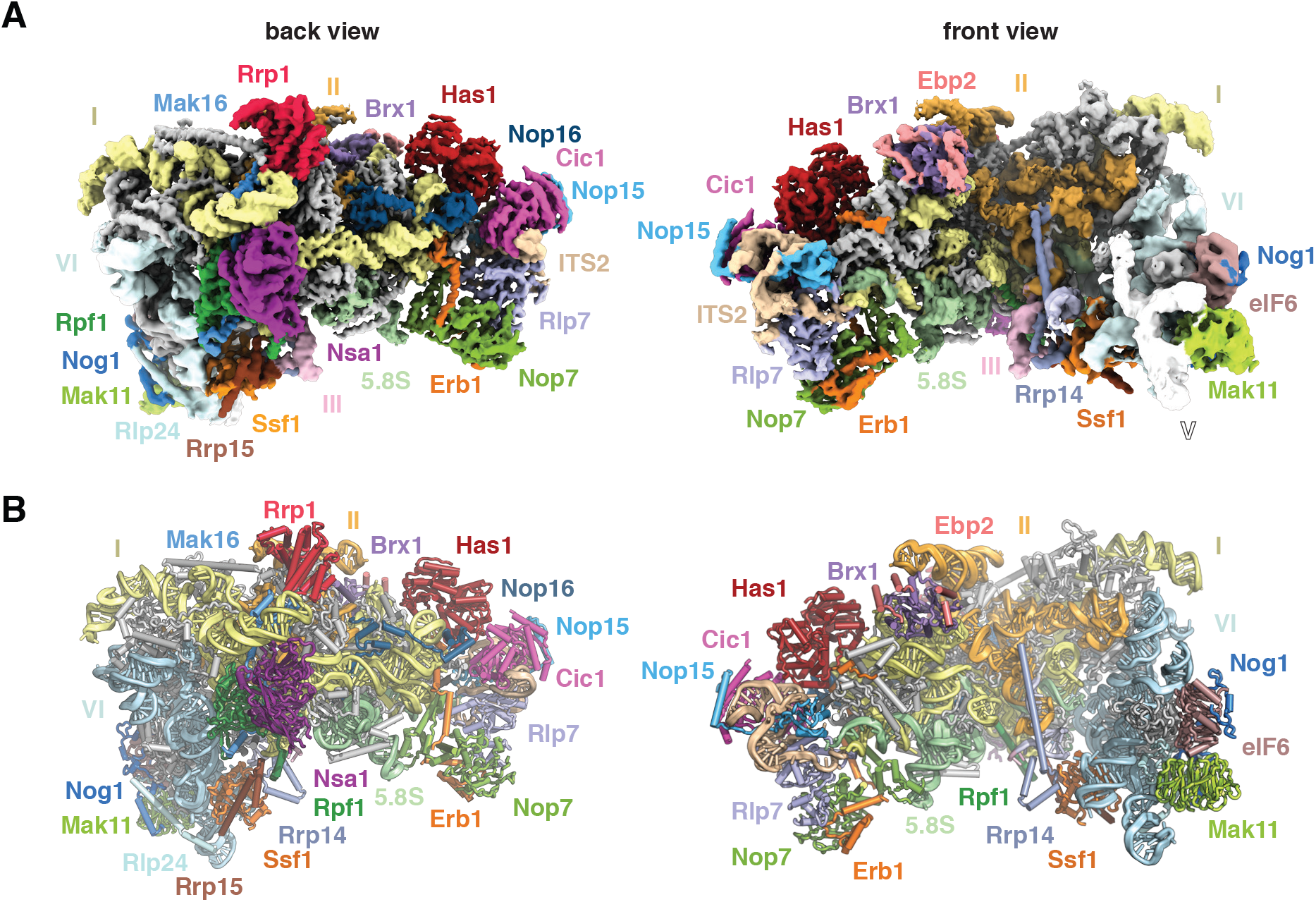
Structure of the early nucleolar large subunit processome. **(A)** Composite cryo-electron density map consisting of 25S rRNA domains I and II (3.4 Å) and VI (3.7 Å, low pass filtered to 5 Å) and associated proteins. **(B)** Corresponding near-atomic model with ribosome assembly factors and 25S rRNA domains labeled and color-coded. Ribosomal proteins are shown in grey.

The purified pre-60S particles contain 27SB rRNA, which has not yet been processed at ITS2 (**fig. S1**). We observed three different conformational states of this RNA (**fig. S2**). State 1 includes ordered density for ITS2 and domains I, II and the 5.8S rRNA. State 2 additionally revealed density for domain VI, which is present in a near-mature conformation. Although present, the majority of domains IV, V and the 5S RNP were poorly resolved in our reconstructions, due to conformational flexibility. Low-resolution features corresponding to parts of domain V (helices 74-79) and its proximal assembly factor Mak11 can be seen in states 2 and 2A. In contrast to state 2, state 3 lacks an ordered domain VI but features domain III.

## Nucleolar Assembly Factors Stabilize rRNA Domain Interfaces

A striking feature of the LSU processome is its open architecture where the solvent-exposed domains I, II and VI are encapsulated by a series of ribosome assembly factors as visualized in state 2 (**Fig. 1**). Interestingly, ribosomal proteins that have been associated with Diamond Blackfan anemia (9) are located at critical inter-rRNA domain interfaces in the structure, suggesting that their architectural roles are especially important during early nucleolar assembly, where defects can trigger the nucleolar stress response (**fig. S5**).

Domains I, II and VI adopt an open conformation that is chaperoned by eight early ribosome assembly factors, which form a ring-like structure at the solvent-exposed side (**Fig. 2A**). In particular, Brix-domain containing factors (Brx1, Rpf1 and Ssf1) act in conjunction with their respective binding partners (Ebp2, Mak16 and Rrp15) to interconnect these junctions and sterically prevent premature RNA-protein and RNA-RNA contacts. Architectural support for the major interface between domains I and II is provided by Rpf1 and its zinc-binding interaction partner Mak16, the helical repeat protein Rrp1 and the beta-propeller Nsa1. Rpf1 and Nsa1 occupy a region near the domain I binding site of Rpl17, while Mak16 and Rrp1 interface predominantly with ribosomal proteins Rpl4 and Rpl32 within domain II (**Fig. 2B**).

**Fig. 2.**
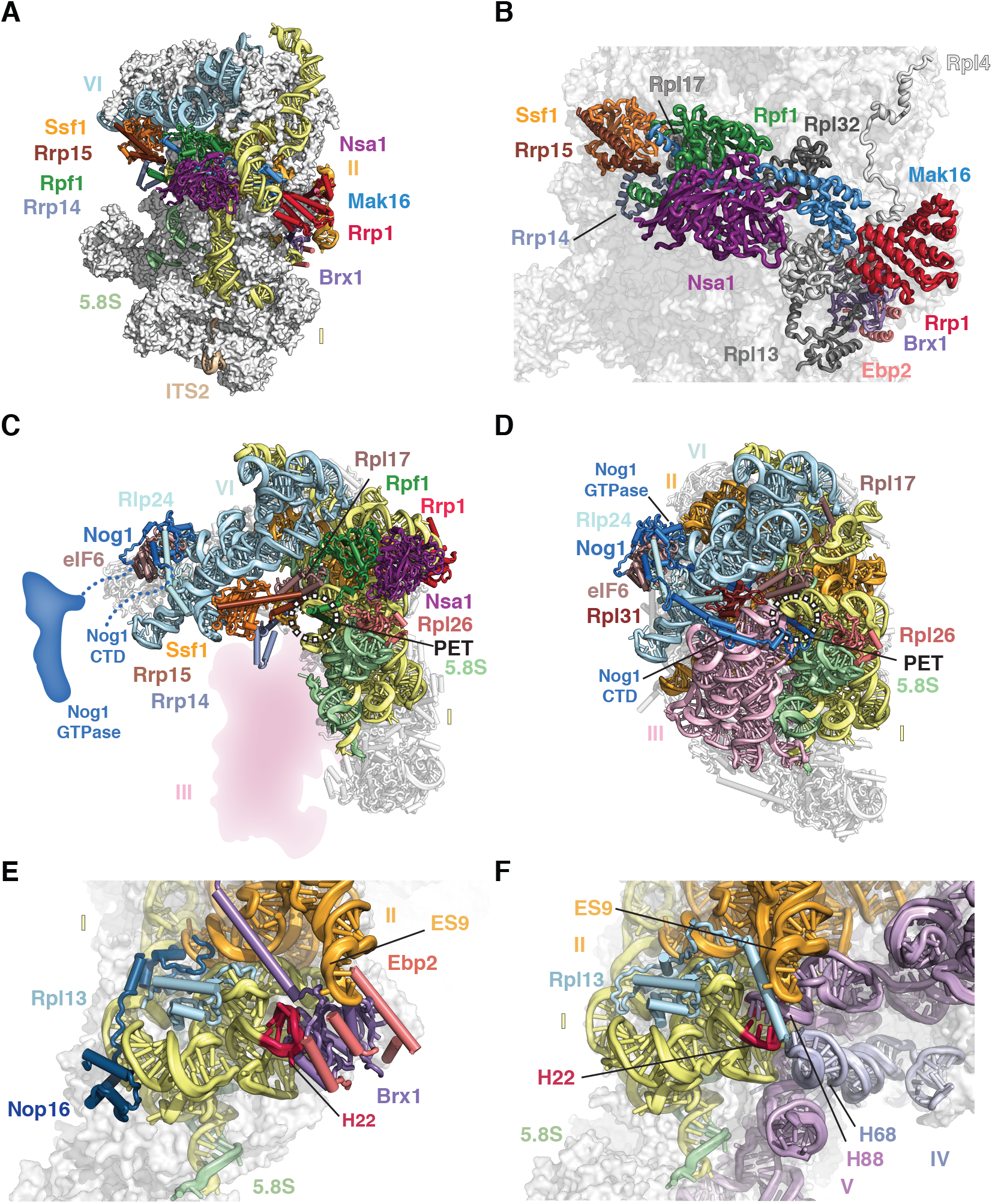
A ring of nucleolar assembly factors prevents premature folding of the 25S rRNA. **(A)** Several assembly factors (colored as in Fig.1) form a ring-like structure to chaperone the solvent exposed areas of domains I, II and VI. **(B)** Interactions of the ring assembly factors with ribosomal proteins Rpl4, Rpl13, Rpl17 and Rpl32 (shades of grey). (**C** and **D**) LSU processome factors prevent the formation of the polypeptide exit tunnel. (C) In the LSU processome (state 2) Ssf1·Rrp15 (orange, brown) block the binding site of Rpl31. Nog1 (blue) is largely unstructured. (D) In the nuclear Nog2-particle (PDB 3JCT), the polypeptide exit tunnel is formed, probed by the Nog1 CTD and supported by Rpl31 (brown). (**E** and **F**) Steric hindrance during nucleolar maturation. (E) Brx1.Ebp2 (purple/pink) remodel domain I (helix 22) to prevent its interaction with domain V (helix 88) while sterically hindering the association of domain IV (helix 68) and the C-terminus of Rpl13 (F). These interactions are visualized in the LSU processome (C and E) and the Nog2-particle (D and F).

The ring-like structure encapsulating domains I, II and VI is continued in one direction by the Ssf1-Rrp15 heterodimer and Rrp14. While the long C-terminal helix of Rrp14 bridges domains II and VI, the Ssf1·-Rrp15 complex is positioned at the interface of domains I and VI (**Fig. 2A**). Here, Ssf1 occupies the same position as Rpl31, which in the later Nog2-containing pre-60S binds at the interface of domains III and VI near the polypeptide exit tunnel (PET) (2). The PET, which is created by domains I, III and VI at the solvent exposed side, is already formed in the Nog2 particle, where it is blocked by the C-terminal domain (CTD) of the GTPase Nog1 (**Fig. 2C** and **2D**).

The role of the Brx1·Ebp2 heterodimer near the interface of domains I and II, is two-fold. While being involved in the stabilization of these domains, its strategic binding site prevents the premature assembly of the large subunit by steric hindrance. In later stages of LSU assembly, an RNA segment of domain I (helix 22) base-pairs with a region in domain V (helix 88) near a separate region of domain IV (helix 68). Brx1 remodels helix 22 of domain I to prevent the premature formation of this tertiary structure with helix 88. Similarly, Brx1 prevents the mature conformations of domain IV (helix 68), domain II (expansion segment 9) and the C-terminal region of Rpl13 in this region (**Fig. 2E** and **F**).

In state 3, we have identified the Erb1·Ytm1 heterodimer bound to domain III via Rpl27 (**Fig. 3A**). The N-terminal region of Erb1 (residues 239-397) wraps around the entire ITS2-domain I interface and is positioned underneath Nop16 and Has1 (**Fig. 3B, C**). This location is in agreement with previous cross-linking data and explains why deletions within this region prevent incorporation into pre-60S particles (10, 11). Nop16 interconnects RNA elements of the 5.8S rRNA and regions of domain I, and additionally contains a bipartite binding site by interacting with ribosomal proteins Rpl8 and Rpl13 (**Fig. 3C**). The DEAD-box helicase Has1 is positioned at the interface of Rpl8, Cic1, Nop16 and Erb1 (**Fig. 3C**). Assembly factors Cic1, Rlp7, Nop7 and Nop15 appear both in the LSU processome and the Nog2-particle in largely the same conformation (**Fig. 3C** and **D**).

**Fig. 3.**
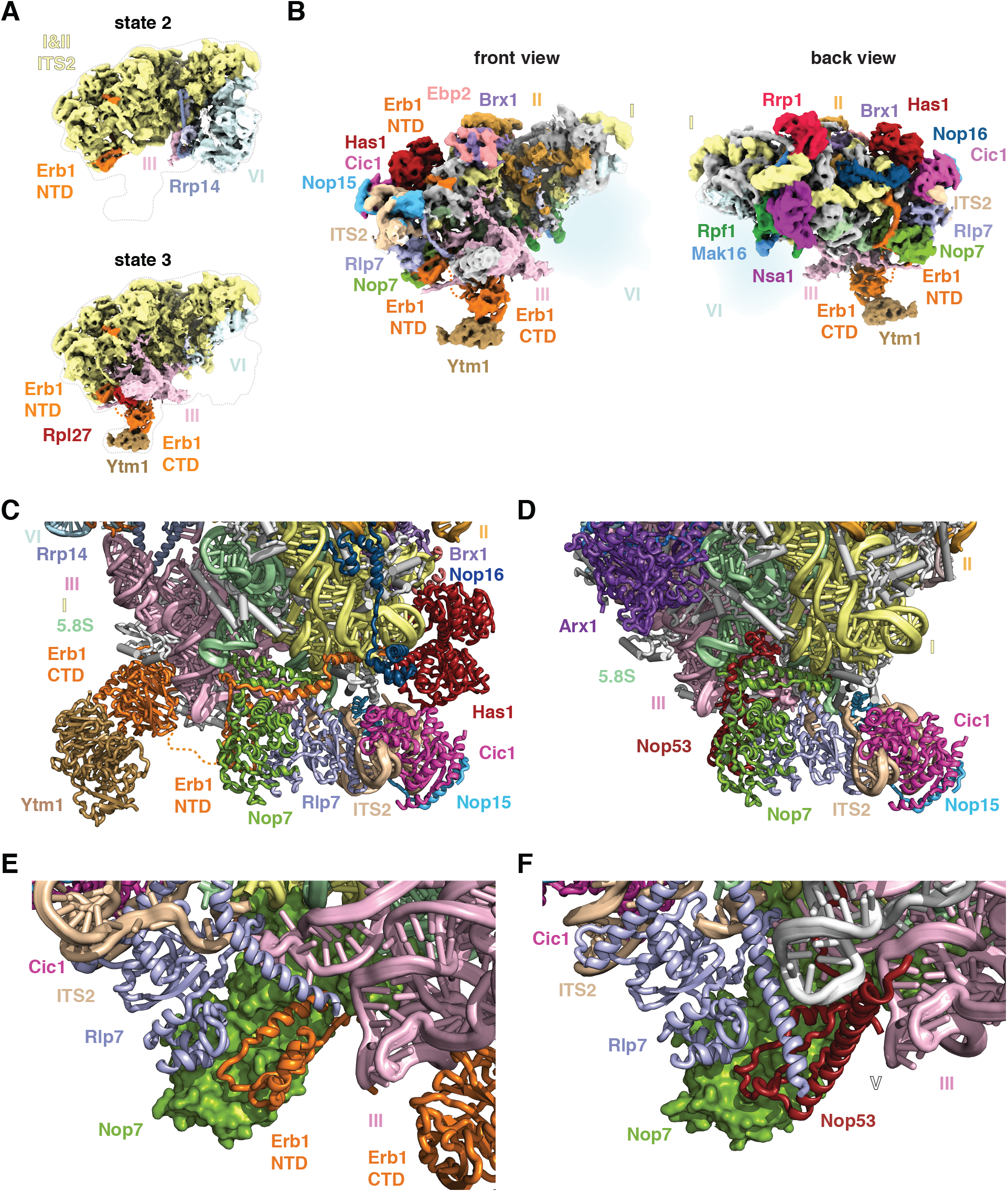
Molecular mimicry by Erb1 prevents premature ITS2 processing. **(A)** Cryo-EM maps of states 2 and 3 of the LSU processome. State 2 contains domain VI (light blue) but limited density for domain III (pink), while state 3 contains domain III and a flexible domain VI. **(B)** Two views of the state 3 cryo-EM map with ribosomal proteins (grey) and color-coded ribosome assembly factors and 25S rRNA domains. (**C** and **D**) Cartoon representation of the ITS2 region in state 3 of the LSU processome (C) and the Nog2-particle (PDB 3JCT) (D). (**E** and **F**) The N-termini of Erb1 and Rlp7 in the LSU processome prevent the binding of Nop53 (E), which binds to Nop7 in the Nog2-particle (F).

Strikingly, the N-terminal segment of Erb1 employs molecular mimicry by binding to Nop7 in a similar fashion as Nop53 in the Nog2-particle, which uses a structurally related motif to bind to Nop7. This steric hindrance is exacerbated by the alternate conformation of the Rlp7 N-terminus that further prevents Nop53 binding (**Fig. 3E** and **F**). Therefore, the coordinated mechanical removal of Erb1 and its proximal factors Ytm1, Nop16 and Has1 by Mdn1 is required before Nop53 can bind to the Nog2-particle and recruit the exosome-associated RNA helicase Mtr4 for ITS2 processing (12, 13). The Has1 helicase may have acted upon its substrate at an earlier stage during the 27SA3 to 27SB transition. Alternatively, it may remodel flexible RNA elements in its vicinity for the ensuing 27SB processing (14).

## Nucleolar Assembly of the Large Ribosomal Subunit

The LSU processome states 2 and 3 represent distinct assembly intermediates of the polypeptide exit tunnel (**Fig. 4**). Ssf1, Rrp15 and Rrp14 are ordered in state 2 where they chaperone domains I and VI, that line two sides of the forming polypeptide exit tunnel (PET). By contrast, domain VI, Ssf1, Rrp15 and Rrp14 are disordered in state 3. Here, Ytm1 and Erb1 chaperone domain III, which adopts a mature conformation with respect to domain I to form a different intermediate of the PET.

**Fig. 4.**
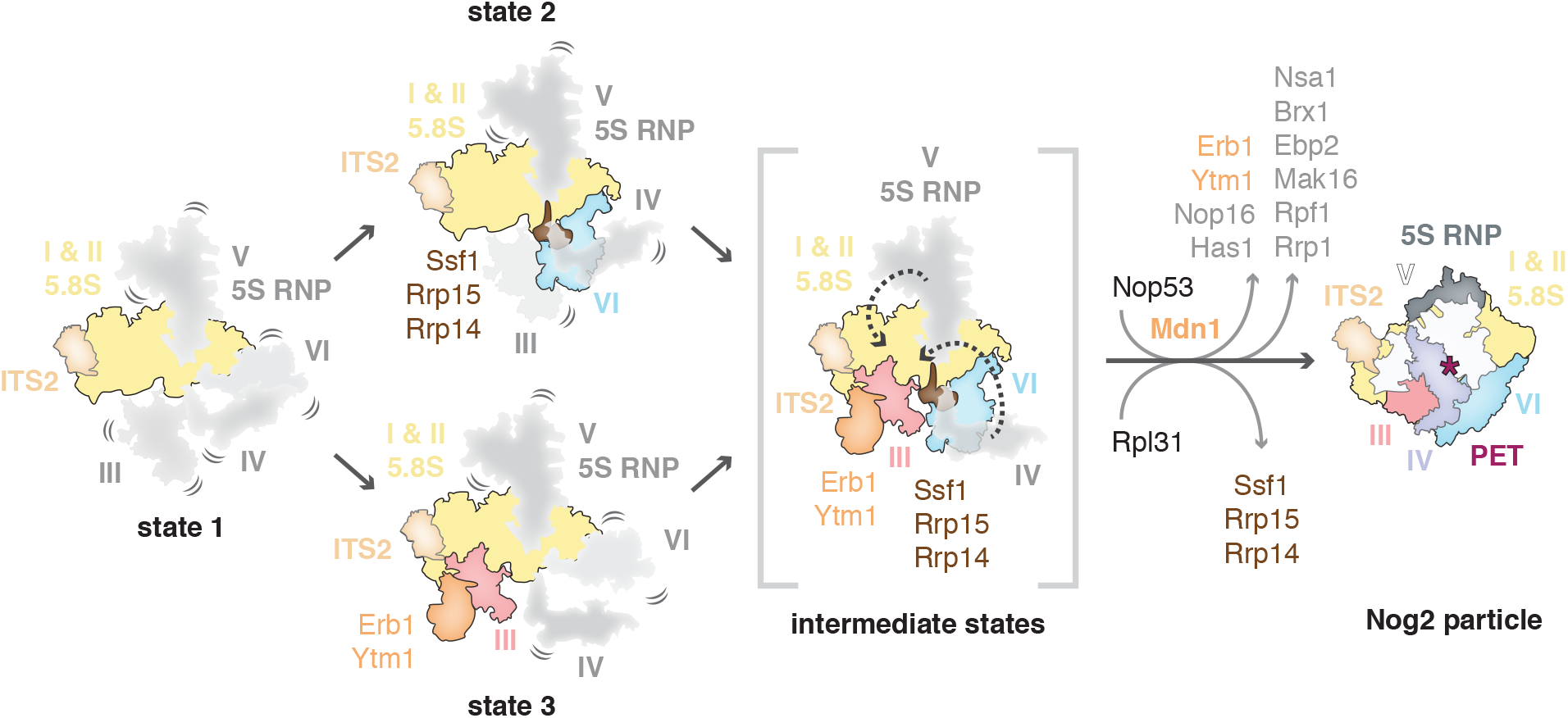
Model for early nucleolar stages of large subunit assembly. Domains of the 25S rRNA are represented as separate segments that are flexible (in grey) or stably incorporated (in color). The LSU processome state 1 represents an intermediate in which subsequent folding of either domain III or VI will occur (state 3 and state 2, respectively) to partially form the polypeptide exit tunnel (PET). To mature into the Nog2-particle, the formation of the nascent polypeptide exit tunnel must occur by Mdn1 and Rix7-dependent exchange and removal of assembly factors alongside the stepwise folding of domain V followed by domain IV.

A subsequent maturation step of states 2 and 3 likely involves the joining of domains III and VI and the formation of the PET on the solvent exposed side. This may be accompanied by the insertion of the Nog1 N-terminus into the nascent PET and the replacement of Ssf1·Rrp15 by Rpl31 (**Figs. 3D** and **4**).

Conformational changes first by the initially flexible domain V/5S RNP together with the Nog1 GTPase domain and subsequently domain IV will result in the overall conformation observed in the late-nucleolar Nog 2-particle (**Figs. 3D** and **4**) (2). The base-pairing between domains I and V will be possible upon the dissociation of the Brx1·Ebp2 complex (**Fig. 2E** and **F**) while the release of Rrp1, Rpf1 and Mak16 is likely brought about by the ATPase Rix7 acting upon the proximal Nsa1 (15). In the ITS2 region, Mdn1-dependent removal of Erb1·Ytm1 will expose the binding site of Nop53 and may also trigger the exit of Nop16 and Has1 from the particle (13).

Nucleolar 60S ribosome assembly is conceptually reminiscent of the early prokaryotic 50S assembly intermediates where different rRNA domains are assembled in a modular fashion (16) (**Fig. 4**). However, the structure of the eukaryotic large subunit processome highlights the high degree of control that is exerted to prevent the premature formation of inter-domain contacts of ribosomal RNA and illustrates how a chronology of assembly factors is enforced. This overarching theme exists for nucleolar stages of both small and large subunit processomes (17).

## Acknowledgements

We thank M. Ebrahim and J. Sotiris for their outstanding support with data collection at the Evelyn Gruss Lipper Cryo-EM resource center, and M. Tesic-Mark for analysis of mass spectrometry data. We further thank the Walz lab for helpful discussions. L.M. is supported in part by NIH T32 GM115327-Tan. J.B. is supported by an EMBO long-term fellowship (ALTF 51-2014) and a Swiss National Science Foundation fellowship (155515). M.C.-M. is supported by a postgraduate scholarship from NSERC. S.K. is supported by the Robertson Foundation, the Irma T. Hirschl Trust, the Alexandrine and Alexander L. Sinsheimer Fund, the Rita Allen Foundation, and an NIH New Innovator Award (1DP2GM123459). B.T.C. is supported by National Institute of Health Grant Nos. P41GM103314 and P41GM109824.

The cryo-EM density map for the yeast LSU processome has been deposited in the EM Data Bank with accession code EMD-XXXX. Atomic coordinates for the yeast LSU processome have been deposited in the Protein Data Bank under accession code XXXX. A PyMOL script for the analysis of the LSU processome will be made available in **Data File S1.**

S.K. and Z.S. established purification conditions. Z.S., L.M., and S.K. determined the cryo-EM structure of the yeast LSU processome. K.R.M., J.W., and B.T.C. processed and analyzed DSS cross-linking data. Z.S., L.M., J.B., M.C.-M.,and S.K. built the model, L.M. performed all RNA work, and Z.S., L.M., J.B., M.H., M.C.-M., and S.K. interpreted the results and wrote the manuscript.

